# GPRIN1 modulates neuronal signal transduction and affects mouse-learning behavior

**DOI:** 10.1101/291377

**Authors:** Claudia Savoia, Julien B Pujol, Angelique Vaucher, Umberto De Marchi, Claus Rieker, Eija Heikkilä, Antonio Núñez Galindo, Loïc Dayon, Elhadji M Dioum

## Abstract

In the adult and developing brain, G protein-regulated inducer of neurite outgrowth 1 (GPRIN1) is a downstream effector for Gα_o/i/z_ proteins that promotes neurite outgrowth. However, so far, the physiological role of GPRIN1 in brain health has not been reported. We generated a viable GPRIN1 whole-body knockout mouse to assess its physiological role in synaptic function both *ex vivo* and *in vivo*. In adult neurons, GPRIN1 is highly localized to the plasma membrane and synapses where it regulates neuronal signal transduction and Ca^2+^ homeostasis. Our results reveal that GPRIN1 might be a novel protein involved in agonist-stimulated cytoskeletal reorganization, which is crucial for early neuronal network development and in functionally mature neurons. Finally, we show that loss of GPRIN1 leads to a learning deficit *in vivo* and sensitizes neurons to stress, suggesting a modulatory role in brain health and disease.

## INTRODUCTION

Neural development is a very dynamic process involving structural remodeling of the soma, neurites outgrowth, axons and dendrites maturation and terminating with synapses generation. Proper synaptic function underlies neuronal activity improving plasticity, learning and memory (Mortimer et al., 2008); (Petros et al., 2008). Receptor-mediated signal transduction is crucial in the regulation of cytoskeleton remodeling through the coordinated signaling of small guanine triphosphate (GTP)ase proteins (or G proteins) (Stiess and Bradke, 2011) (van Kesteren and Spencer, 2003) (Flavell and Greenberg, 2008; Mattson, 2008). Ligand-dependent activation of Gα_o/i/z_-protein-coupled receptors has been implicated in the regulation of neurite outgrowth and neuronal differentiation (Ge et al., 2009; Lotto et al., 1999; Reinoso et al., 1996). In an attempt to identify novel effectors of activated Gα_o/i/z_ proteins, the G protein-regulated inducer of neurite outgrowth (GPRIN1) was discovered by Chen and colleagues as a membrane-bound protein enriched in the growth cones of neurites and an activator of neurite outgrowth (Chen et al., 1999). Subsequently, several reports proposed a putative role for GPRIN1 in the adult and developing brain as a downstream effector for Gα protein signaling, mainly due to its specific and preferential interaction with the activated Gα_0_ and to the presence of conserved common motifs important for Gα_0_ and for Sprouty2 binding. While its interaction with Sprouty2 has been reported to modulate the MAPK pathway (Hwangpo et al., 2012), GPRIN1-GTP-Gα_0_ interaction has been found to be essential to recruit potential partners involved in cytoskeleton reorganization and receptor localization (Nakata and Kozasa, 2005) and to affect neurite outgrowth directly regulating µ-Opioid receptor activation within the membrane lipid raft (Ge et al., 2009). Recently, other neuronal receptors, like nicotinic acetylcholine receptor (AchR) a7, a4 and b2 subunits, have been coupled to a G protein complex consisting of GPRIN1, GAP-43 and Cdc42 modulating calcium signaling and axonal development in primary hippocampal neurons (Kabbani et al., 2013; Nordman and Kabbani, 2012; Nordman et al., 2014). Furthermore, GPRIN1 is phosphorylated by Cyclin-dependent kinase 5 (Cdk5), an essential protein known to regulate cytoskeleton remodeling, axonal guidance and neuronal plasticity in the brain (Contreras-Vallejos et al., 2014; Kang et al., 2008). GPRIN1 expression is partially regulated by promoter recruitment of the methyl-CpG binding protein 2 (MeCP2), a transcriptional regulator mutated in a plethora of neurobehavioral abnormalities ranging from learning difficulties to autism-like behaviors in Rett syndrome (Chahrour et al., 2008)(Ito-Ishida et al., 2015). These observations further confirmed the unexplored importance of GPRIN1 and the need to investigate its physiological role in neuronal dynamics and function. To this aim, we generated for the first time a viable mouse lacking whole-body expression of GPRIN1, and we assessed the role of GPRIN1 in synaptic function both *ex vivo* and *in vivo*. Here, we showed that GPRIN1 was strongly expressed in adult mouse brain, enriched in the hippocampus, hypothalamus and habenula, as well as throughout the cerebral cortex. Our results demonstrate that GPRIN1 plays a critical role not only in early stages of neuronal network development, but also in functionally mature neurons affecting calcium signaling after agonist-induced physiological response and contributing to spontaneous neuronal electrical activity. Furthermore, loss of GPRIN1 sensitized primary neurons to stress *ex vivo*, further supporting the learning defects observed in adult GPRIN1^−/−^ mice. In this study, we propose GPRIN1 as a novel regulatory protein involved in physiological neuronal function and learning behavior, opening further questions related to its role in brain health and neurodegeneration.

## EXPERIMENTAL PROCEDURES

### Primary neuronal cultures

Cortical neurons from GPRIN1^−/−^ mice and WT littermates were prepared from E16.5 embryos by using a previously described method (Abramov et al., 2007) with few modifications. Briefly, mice were anesthetized and euthanised. Embryonic brain tissue was dissociated by trituration in Hanks’ Balanced Salt Solution (HBSS) supplemented with Hepes (10mM), then trypsinized (0.025%) for 5 min at 37 °C. After centrifugation, cell suspension was plated on Poly-L-ornithine (PLO)-coated coverslips (20 μg/ml, Sigma) and maintained in Neurobasal medium (NB) (Invitrogen) supplemented with B-27 (Invitrogen), 0.5 mM L-glutamine and 2-mercaptoethanol (50 μM). Cultures were maintained at 37 °C in a humidified atmosphere of 5% CO_2_ and 95% air, fed once every 3 days and maintained for a minimum of 5 days before experimental use. Cytosine arabinoside (Ara-C, 5 μM) was added within 48 hours from plating to prevent the growth of non-neuronal cells.

### Cell culture

SH-SY5Y neuroblastoma cells were obtained from the American Tissue Culture Collection (ATCC). Cells were cultured in Dulbecco’s modified Eagle’s medium (DMEM, Invitrogen) containing glucose, l-glutamine and sodium pyruvate (1 mM). This medium was supplemented with 10% (v/v) heat-inactivated fetal bovine serum (FBS, Invitrogen) and 1% penicillin streptomycin (P/S, Invitrogen). To knock down endogenous human Gprin1 we infected cells with shRNA targeting Gprin1 using adenoviral vector generated by Sirion Biotech (Martinsried, Germany) following manufacturer’s instructions.

### Reagents

All the treatments were added to B27-free NB or HBSS modified with 10 mM HEPES without phenol red (Invitrogen). (s)-a-Amino-3-hydroxy-5-methyl-4-isoxazolepropionic acid ((S)AMPA) and N-methyl-D-aspartic acid (NMDA) were obtained from Tocris Bioscience (Bristol, UK) and dissolved in water and used at a final [50-200 μM]. 8-(4-Chlorophenyl)thio-cyclic AMP (8-CPT-cAMP) was dissolved in water and used at final [0.5 mM]. Forskolin (FSK) and IBMX were dissolved in DMSO and used at final [10μ] and [100 nM], respectively. Cyclohexamide (CHX), TNF alpha, H_2_O_2_ (Sigma) and KCl were used at final [10 μ/mL], [10 ng/mL], [200 μM] and [30 mM], respectively. Custom sheep antibody against full-length GPRIN1 protein was generated by MRC PPU Reagents and Services, University of Dundee, UK.

### Multi-Electrode Array (MEA)

Spontaneous electrical activity of primary cortical and hippocampal neurons was measured using the MEA system (Axion BioSystems Maestro). Neurons were plated on PLO-coated MEA 96-or 12-well plates (140,000 neurons/well). Baseline electrical activity was recorded for 20 min on a heated stage at 37°C every day *in vitro* (DIV) in order to have the timeline activity data set. Half of the medium was replaced every 3 days with fresh complete medium. Treatments were applied starting from DIV 14. Briefly, 100 μL of conditioned media was removed from the MEA plate and reagents in 100 μL NB fresh medium were slowly added back to the cells. The MEA plate was returned to the stage and recordings resumed immediately, and lasted for at least 30 min (network dynamics). To control for possible solvent effects as well as mechanical artifacts arising from the exchange of solutions, an additional column treated with DMSO was recorded (vehicle). Data analysis was performed using Axion Integrated Studio (AxIS) software after low-frequency components removal by high-pass filtering all traces at 200 Hz. The first 10 ms after the stimulus were discarded in order to avoid any possible effect of the stimulus artifact. Spike and Burst Detector data processors were used to analyze spike timing patterns to identify bursts on individual electrodes (single-electrode bursts) and across multiple electrodes (network bursts).

### Single-cell imaging of cytosolic calcium signals

Intracellular calcium influx [Ca^2+^]i was measured in primary neurons as described previously (Barreto-Chang and Dolmetsch, 2009). Briefly, primary neurons were plated on PLO-treated 35-mm-diameter glass-bottom dishes (MatTek). After one week in culture, neurons were washed twice with Krebs-Ringer bicarbonate Hepes buffer (KRBH) containing 140 mM NaCl, 3.6 mM KCl, 0.5 mM, NaH2PO4, 0.5 mM MgSO4, 1.5 mM CaCl2, 10 mM Hepes, and 5mM NaHCO3 (pH 7.4) and loaded with 2.5 μM Fura-2AM (Life Technologies) for 30 min at room temperature (RT) in the dark. The neurons were washed twice with KRBH and then incubated for 30 min to allow complete de-esterification of intracellular Fura-2AM ester. Glass coverslips were mounted on a DMI6000 B inverted fluorescence microscope using an HCXPL APO 40 oil-immersion objective (Leica Microsystems) and an Evolve 512 back-illuminated CCD with 16 pixels camera (Photometrics, Tucson, AZ). Images were taken every 2 sec, and the fluorescence ratios (F340/F380) representing the corresponding [Ca^2+^]i changes were calculated using MetaFluor 7.0 (Meta Imaging Series) and analyzed in Microsoft Excel and GraphPad Prism 5.

### Imaging

For imaging primary neurons, 1×10^5^ cells were seeded onto PLO-coated 13 mm round glass coverslips. Fixed cells were immunostained with primary antibodies (details in **Supplementary Table 1**) and isotype-matched AlexaFluor-conjugated secondary antibody (Molecular Probes). Images were acquired for each channel using a Leica SP8 microscope with a 63×1.4 NA Plan Apochromat objective (Leica) and processed using ImageJ (NIH). Quantitative analysis of total neurite length was performed using the NeuronJ plugin for ImageJ (Meijering et al., 2004).

For live imaging experiments, primary neurons seeded on PLO-coated 15 mm MatTek coverslips were infected with GPRIN1-GFP after 3 DIV using adenoviral vector generated by Sirion Biotech (Martinsried, Germany), following manufacturer’s instructions. 48 hours post-infection, cells were incubated with Sir-Actin (Spirochrome, Cytoskeleton, Inc) to visualize cytoskeleton, and live time-lapse imaging was performed on a Leica SP8 microscope equipped with a CO_2_/temperature-controlled system and a 63×1.4 NA Plan Apochromat objective (Leica).

### Automated high-image content acquisition and analysis

For high-content imaging, primary neurons were fixed in 4% PFA for 15 min at RT. Primary antibodies diluted in blocking buffer were applied overnight (ON) at 4 °C. Dapi (Invitrogen) and secondary antibodies conjugated to Alexa fluorophores (Molecular Probes) were diluted at 1:10,000 in blocking buffer and applied for 2 hours at RT. To quantify synapses, image acquisition and analyses were performed using the ImageXpress Micro XLS system (Molecular Devices, Sunnyvale, CA, USA). A presynaptic marker (anti-synaptophysin antibody; 1:500 ref. ab32127) and a postsynaptic marker (anti-PSD-95 antibody; 1:300; ref. MA1-046) were used and multiplexed with the neuronal marker (anti-beta III Tubulin, Tuj-1; 1:1,000; ref. ab107216) to identify neurons and create a “neurite” mask. An algorithm was then generated using MetaXpress™ software application modules in order to identify pre- and post-synaptic spots specifically located on neurites, and then to count the number of spot co-localizations (considered as synapses). Results were expressed as the density of spots for 100 μm of neurite. A minimum of 10 sites per well (approximately 2,000 neurons) was analyzed.

### Mitochondrial Respiration Assay

Oxygen consumption rate (OCR) in live intact neurons was measured using the Seahorse XF Extracellular Flux 96 analyzer according to the manufacturer’s instructions (Seahorse Bioscience) (Ribeiro et al., 2015). In brief, neurons were seeded into POL-coated Seahorse tissue culture plates (25,000 cells per well) and sequentially exposed to control assay medium, oligomycin (ATP synthase inhibitor), receptor agonists, and a mix of rotenone (complex I inhibitor) and antimycin A (complex III inhibitor). Following the serially injection of these compounds, OCR was used to calculate different domains of mitochondrial function such as baseline respiration, ATP turnover, proton leak respiration and non-mitochondrial respiration (**Fig. S4A**). OCR data were normalized to protein concentration on a per well basis.

### Release of endogenous BDNF by ELISA

BDNF in culture supernatants was measured by using the mouse BDNF PicoKine™ ELISA Kit (Boster, Wuhan, CHN), according to the manufacturer’s protocols. Briefly, medium was collected after stimulation with (S)AMPA and added to a plate pre-coated with BDNF specific antibody and incubated at 37 °C for 90 min. Then a biotinylated anti-mouse BDNF antibody was incubated at 37 °C for 60 min, followed by addition of avidin-biotin-peroxidase-complex for 30 min at 37 °C and color development. Absorbance was measured at 450 nm on a multi-mode microplate reader (BioTek Instruments, INC., USA). The BDNF assay had a linear over a range of 0-31.2 pg/ml and BDNF levels were adjusted to the amount of protein in the corresponding culture well (pg BDNF/mg protein) (Xie et al., 2017).

### Western Blotting and Antibodies

Protein samples (15-20 µg) were loaded in 4–12% Tris–MOPS-SDS–PAGE (NuPage, Invitrogen) after BCA protein quantification (Bio-Rad). The proteins were transferred from the gel onto nitrocellulose membranes (Bio-Rad) by wet transfer. The blot was blocked for 1 hour at RT in a 1:1 Li-Cor blocking buffer: 1X TBS plus 0.05% Tween and incubated ON at 4° C with the primary antibodies (details in **Supplementary Table 1**). Fluorescent labeled Li-Cor secondary antibodies were incubated for 1 hour at RT. After several washes in TBS-T, blots were scanned with a Li-Cor Odyssey.

### RNA isolation and qPCR

Total RNA was prepared by RNA mini kit Plus (QIAGEN). RT-PCR was performed with the Applied High Capacity cDNA Synthesis kit (Thermo Scientific) and cDNA were used for qPCR analysis. The target gene expression was evaluated using Power SYBR Green PCR Master Mix (Applied Biosystems). PCR was carried out on a LightCycler 480 Real-Time PCR Systems (Roche) using a LightCycler 1536 SYBR green (Roche). The thermocycling conditions used were as follows: an initial step of 10 min at +95 °C, 45 cycles of a 10 sec denaturation at +95 °C, 10 sec annealing at 60 °C, and an 11 sec extension at +72 °C. Transcript levels were normalized to cyclophilin B. Relative fold change in expression was calculated using the ΔCT method. For relative transcript quantification, each cDNA sample was run on a 4-point standard curve so as to assure a PCR efficiency of ≥95%. Primer sequences are listed in **Supplementary Table 2**.

### Animal experimentation

This study was performed in strict compliance with the local guidelines for the care and use of laboratory animals. Prior approval by the local ethical committees was obtained for all protocols. The experimental group was made of 24 mice 9-12 weeks old (6 males and 6 females GPRIN1^−/−^ mice + 6 males and 6 females littermates WT mice).

| Pipeline study | Date |
| --- | --- |
| Habituation | Week 1 |
| Open field | Week 2 |
| Water maze | Week 3 |
| Fear conditioning | Week 4 |

### Morris Water Maze

The Morris Water Maze (MWM) task was used to assess spatial learning (Morris, 1984). Briefly, mice had to learn to find a hidden platform in a circular water tank using visual cues. First, mice were tested for learning during 4 training days, with 6 trials per day (120 sec each with inter-trial interval of 30 min). At the beginning of a trial, the mouse was placed in the water facing the edge of the tank at one of the 4 starting points (North-East, South-East, South-West, and North-West). The starting points were changed from trial to trial, but the escape platform was fixed (West quadrant) throughout the experiment. Only if the mouse was able to find the platform within 120 sec was it allowed to remain, otherwise the trial was stopped. After training, the platform was removed from the tank and a probe test with 1 trial of 120 sec was performed. Latencies to find the escape platform, radius around the escape platform, time and distance traveled in the target quadrant were recorded and analyzed in GraphPad Prism 5.

### Fear Conditioning

The Fear Conditioning task was performed to assess emotional learning and memory. On the first day (Training), the animals were placed in a metal sidewalls and stainless-steel grid. After 2 min of habituation to the chamber, the tone-foot shock pairing (tone: 5000 Hz, 80 dB, 30 sec; shock: 0.7 mA, 2 sec) was presented and repeated 4 times with a 2-min interval. On the second day (Context test), no tone and no shock were presented to the mouse. The following day, the final test (Tone) was assessed, moving the mice to a curved wall in white PVC plate, and after 2 min of habituation, the tone (5000 Hz, 80 dB) was presented continuously for 8 min. During each session (Training: day 1, Context: day 2, Tone: day 3), the percentage of the total duration of freezing expressed as “time freezing” was recorded and analyzed in GraphPad Prism 5.

### Brain Histology

The whole brain was dissected from a previously euthanized mouse. The brains were removed and post-fixed in 4% paraformaldehyde in PBS overnight (ON) at 4 °C. Tissue was cryo-protected in 20% sucrose in PBS at 4 °C, then frozen with dry ice in disposable plastic Cryomold covered with OCT (optimum cutting temperature) compound (Sakura Finetek). 30-μm-thick coronal sections were cut on freezing microtome and stored at −20 °C until use. Frozen brain slices were incubated with 5% normal donkey serum (NDS, Jackson ImmunoResearch Laboratories, West Grove, PA) in PBS with 0.2% Triton-X100 (Sigma-Aldrich, St. Louis, MO). After washing in PBS, slices were incubated ON with primary antibodies (details in **Supplementary Table 1**) at 4 °C, followed by isotype-matched AlexaFluor-conjugated secondary antibody (Molecular Probes). Images were acquired for each channel using a Leica SP8 microscope with a 63×1.4 NA Plan Apochromat objective (Leica) and processed using ImageJ (NIH). Cresyl Violet (CV) staining was performed using the fully automated Ventana Discovery XT (Roche Diagnostics, Rotkreuz, Switzerland).

### Statistical analysis

Statistics were performed with GraphPad Prism 5 (GraphPad Software, San Diego, CA, USA). Data are represented as mean ± SEM of 3 independent experiments unless otherwise indicated. Data were analyzed by the Student’s t-test for paired observations. When three or more means were compared, ANOVA was applied followed by multiple comparison tests. Value of *p<0.05 was considered statistically significant.

## RESULTS

### *In vivo* and e*x vivo* characterization of genetic GPRIN1-ablated neurons

GPRIN1 has been previously associated with neurite outgrowth in neuronal cell lines, however its physiological function in post-mitotic neurons has not been described yet. We therefore set out to investigate the consequences of neuronal GPRIN1 ablation by combining *ex vivo* and *in vivo* models of GPRIN1 deletion. The viable GPRIN1 whole-body knockout (GPRIN1^−/−^) was generated by germline deletion of the floxed GPRIN1 allele after crossing with a Rosa 26-Cre mouse. The GPRIN1^−/−^ mouse did not show defects in general brain architecture and hippocampal structure visualized with Cresyl Violet (CV) staining (**Fig. 1A-S1B**). Histological analysis using a custom antibody against full-length GPRIN1 protein on sagittal adult brain showed that GPRIN1 was strongly enriched in hippocampus, hypothalamus, habenula as well as throughout the cerebral cortex in adult mouse brain (**Fig. S1A-CD**). To gain deeper understanding of the role of GPRIN1 in neuronal function, we used primary cortical and hippocampal neurons isolated from embryos (E16-E18) and kept in culture for several days *in vitro* (DIV). qPCR analysis of neurons after 5, 7 and 11 DIV confirmed the loss of GPRIN1 transcript in the GPRIN1^−/−^ and showed that Gprin1 mRNA expression was higher in the early stage of neuronal differentiation in WT (**Fig. 1D**). Nevertheless, in GPRIN1-ablated neurons, cell viability was unaffected over time compared to WT controls, and cultured cells developed well-interconnected neurite networks (**Fig. 1E-C**). Immunofluorescence (IF) studies displayed GPRIN1 expression mainly at the membranes, along neurites and enriched in growth cones in Tuj1-positive neurons, where it co-localized with GAP43 but not with glia marker S100A (**Fig. 1BC-S1EF**). Interestingly, both GPRIN1 expression and cellular localization pattern were confirmed in human iPSC-derived (iCell) neurons (**Fig. S2AD**), suggesting that this neuronal-abundant protein might play an appealing and still largely unexplored physiological role also in human neurons.

**Figure 1:**
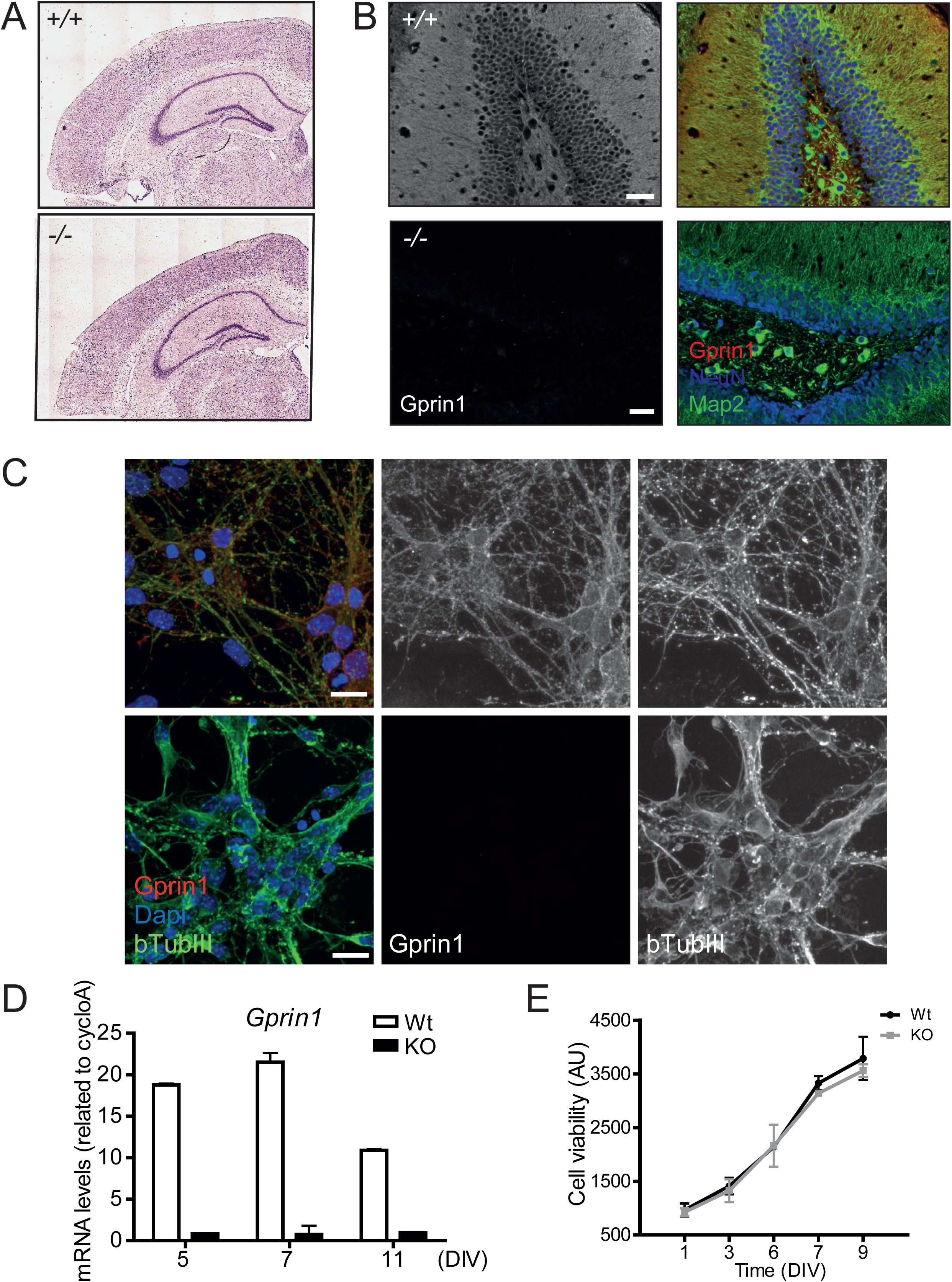
Endogenous GPRIN1 expression and localization in brain and neurons. (A) Histological analysis was performed on sagittal brain slices from adult GPRIN1 WT and GPRIN1^−/−^ mice stained with HE. (B) Immunohistochemistry (IHC) performed on fixed sagittal brain slices from adult GPRIN1 WT and GPRIN1^−/−^ mice stained with Dapi (Invitrogen), GPRIN1 and specific neuronal marker (Map2). (C) Primary cortical neurons from GPRIN1 WT and GPRIN1^−/−^ E16 mice were fixed and stained with Dapi (Invitrogen), GPRIN1 and specific neuronal marker (bTubIII). (D) Quantitative PCR (qPCR) analysis of GPRIN1 mRNA expression in GPRIN1 WT and GPRIN1^−/−^ primary cortical neurons cultured for 5, 7 and 11 days *in vitro* (DIV). (E) Cell viability was assessed over time in GPRIN1 WT and GPRIN1^−/−^ primary neurons cultured on 96 wells at every DIV using Alamar Blue reagent according to the manufacturer’s instructions (Invitrogen). Data were expressed as arbitrary units and represent mean ±SEM from 3 independent experiments.

### Functional analysis of GPRIN1^−/−^ primary cortical neurons

To assess the role played by GPRIN1 during neurite extension, primary neurons were cultured and analyzed over time to visualize morphological differentiation features from isolation towards final maturation. Each differentiation stage has distinguishing characteristics resulting from a highly dynamic cytoskeleton remodeling regulating neurite outgrowth (Dotti et al., 1988). Surprisingly, our results showed that at an early stage *in vitro* (DIV2-3) GPRIN1^−/−^ neurons developed longer neurite processes compared to WT controls; however, at later stages (DIV 5), neurite length was comparable between the two genotypes (**Fig. 2AB**), confirming the already-proposed role of GPRIN1 as gate-keeper of neurite outgrowth. However, our primary goal was to investigate whether GPRIN1 affects neuronal function in mature neurons, since the dynamic changes in morphology are not only required to maintain proper dendrites and branching outgrowth, but mainly allow neuronal maturation and function through spine density increase. Thus, spontaneous electrical activity was recorded by Multi-Electrode Array (MEA), a method that enables simultaneous and long-term recordings of local field potentials and extra-cellular action potentials from a neuronal population at millisecond time-scale (Obien et al., 2014). Our results showed that both GPRIN1 WT and GPRIN1^−/−^ neurons in culture displayed synchronized firing (**Fig. 2C**), and the number of spikes and bursts increased over time across their networks: phenomena directly associated with an increase of synapses density (Ito et al., 2013). Interestingly, GPRIN1^−/−^ neurons showed a significantly reduced number of spikes and bursts (**Fig. 2DE**) compared to WT, indicating that GPRIN1 directly affected basal neuronal activity. To determine whether structural or synaptic abnormalities could cause the functional impairment in the network connectivity observed in GPRIN1^−/−^ neurons, we performed an automated high-content imaging (HCI) of immunostained cultured neurons. Our findings revealed that the amount of pre-synaptic Synapsin (**Fig. 2F**) and post-synaptic density component PSD-95 (**Fig. 2G**) proteins, as well as the number of synaptic puncta (**Fig. 2H**), was comparable between WT and GPRIN1^−/−^ neurons. These results indicate that GPRIN1 is dispensable for brain development and synaptic formation, however, it might be inherently linked with neuronal function.

**Figure 2:**
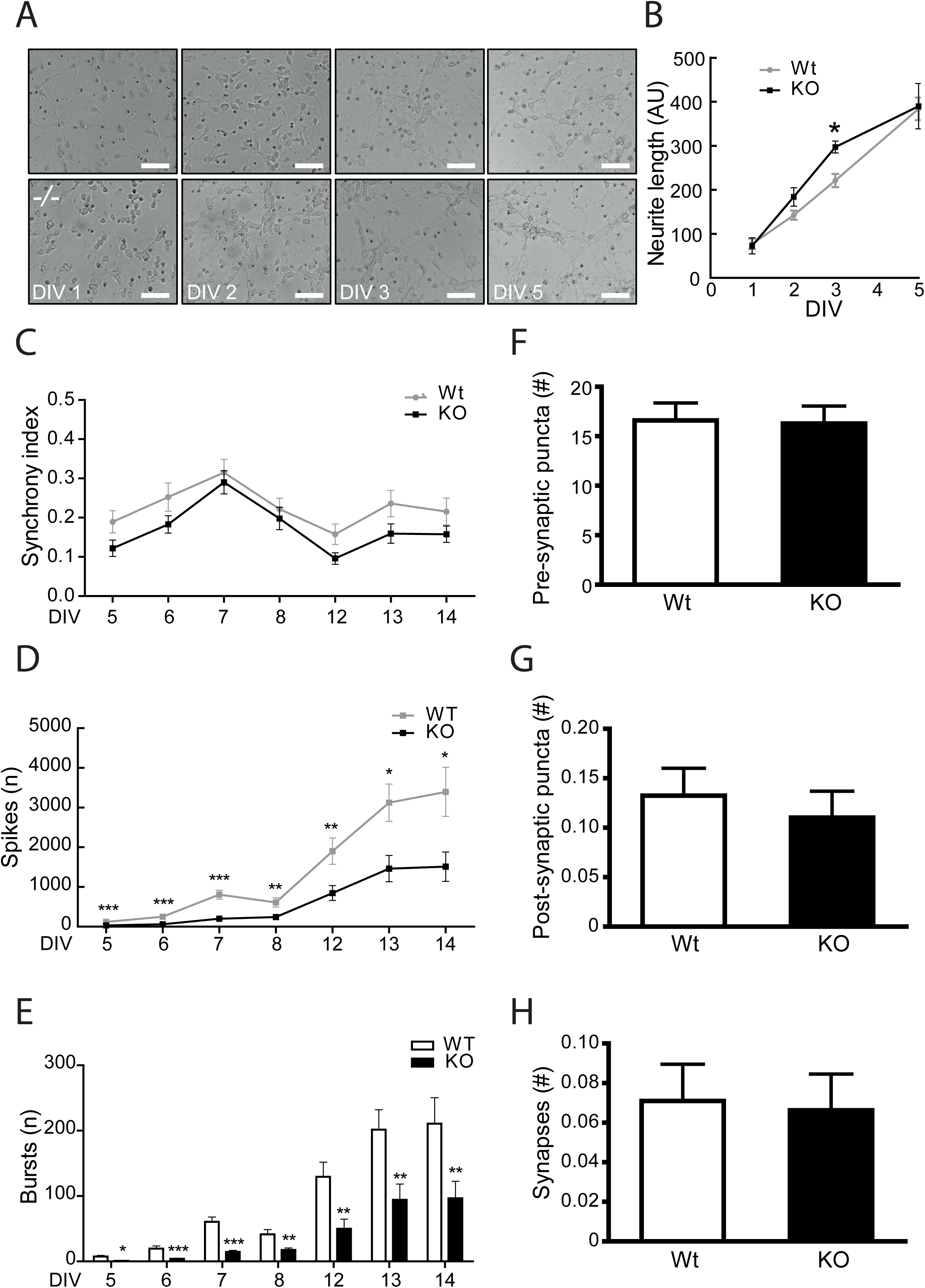
GPRIN1^−/−^ primary neuronal culture characterization. (A) To depict neurites extensions, phase contrast images of GPRIN1 WT and GPRIN1^−/−^ primary neurons were acquired over time, and (B) quantification of neurite length was performed using ImageJ. Data are mean+SEM of 3 independent experiments (*p<0.05) and expressed as arbitrary units (AU). (C) Spontaneous electrical activity of primary neuronal cultures grown up to DIV14 was assessed recording synchrony index, (D) number of spikes and (E) number of bursts. (F) Primary neurons were fixed and labeled with Synapsin (pre-synaptic) and PSD-95 (post-synaptic) markers. Confocal images corresponding to the masks created in each channel were analyzed to quantify pre-, (G) post-synaptic puncta and (H) co-localized puncta corresponding to structural synapses. Results were expressed as mean ± SEM of 3 independent neuronal cultures.

### GPRIN1 effect on calcium homeostasis in physiologically activated neurons

An alternative approach for measuring neuronal activity was to monitor the rapid fluctuations of intracellular calcium (_int_Ca^2+^) levels that accompany neuronal depolarization after physiological activation (Smetters et al., 1999). Indeed, in neuronal cells, Ca^2+^ homeostasis is tightly regulated by several checkpoints, such as G protein-coupled receptors, ion channels, Ca^2+^ binding proteins, transcriptional networks and ion exchangers (De Marchi et al., 2014) to maintain the network integrity and ensure proper neuronal function. Primary neuronal cultures at DIV9 develop spontaneous _int_Ca^2+^ oscillations due to the activity of glutamate, GABA receptors and L-type calcium channels (Dravid and Murray, 2004; Pacico and Mingorance-Le Meur, 2014); (Wang and Gruenstein, 1997). We measured global Ca^2+^ responses by single-cell imaging and found that the number of spontaneous calcium transients, here named as basal mini-peaks, was reduced in GPRIN1^−/−^ compared to WT neurons (**Fig. 3C**). Stimulation with non-toxic concentrations (50-100 μM) of a specific AMPA receptor agonist, (S)AMPA, well-known to activate calcium-dependent exocytosis of pre-synaptic vesicles (Diaz-Trelles et al., 1999; Grienberger and Konnerth, 2012) led to a fast and sharp rise of cytosolic Ca^2+^ in WT (**Fig. 3A**) but not in GPRIN1^−/−^ cortical neurons (**Fig. 3B**). Moreover, the amplitude of the (S)AMPA-induced peak (Max Peak) was significantly reduced in GPRIN1^−/−^ compared to WT neurons (**Fig. 3E**). Interestingly, we observed similar results in neurons stimulated with Carbachol (Cch), an acetylcholine/muscarinic receptor agonist (Mathes and Thompson, 1994) (**Fig. S3A**). Nevertheless, the total amount of Ca^2+^ mobilized after (S)AMPA stimulation did not vary (**Fig. 3F**), although the number of (S)AMPA-evoked electrical spikes was significantly reduced in neurons lacking GPRIN1 (**Fig. 3G**). These observations confirm that GPRIN1 contributes to Ca^2+^ handling, but suggest its involvement only in the first phase of neuronal receptor activation. Furthermore intracellular calcium homeostasis has been linked to mitochondrial function and ATP production, crucial events responsible for sustaining neuronal electrical activity (Rueda et al., 2014). Given these premises we explored the role of GPRIN1 on mitochondrial function of neurons exposed to physiological concentration of AMPA receptor agonist by extracellular flux analysis with Seahorse XF-96 system (Salabei et al., 2014) (Brand and Nicholls, 2011). Mitochondrial respiratory capacity was assessed by sequentially injecting (S)AMPA, oligomycin and a mixture of rotenone and antimycin A (Dranka et al., 2011) to measure basal and stimulated respiration, ATP production, proton leak and non-mitochondrial respiration (**Fig.S4A**). Our results showed that GPRIN1^−/−^ neurons exhibited distinct metabolic profiles with a significant reduction in baseline respiration and ATP turnover (**Fig.3G-I; Fig.S4B-D**). This effect was specific to mitochondrial metabolism as non-mitochondrial respiration and proton leak did not change (**Fig.3JK**).

**Figure 3:**
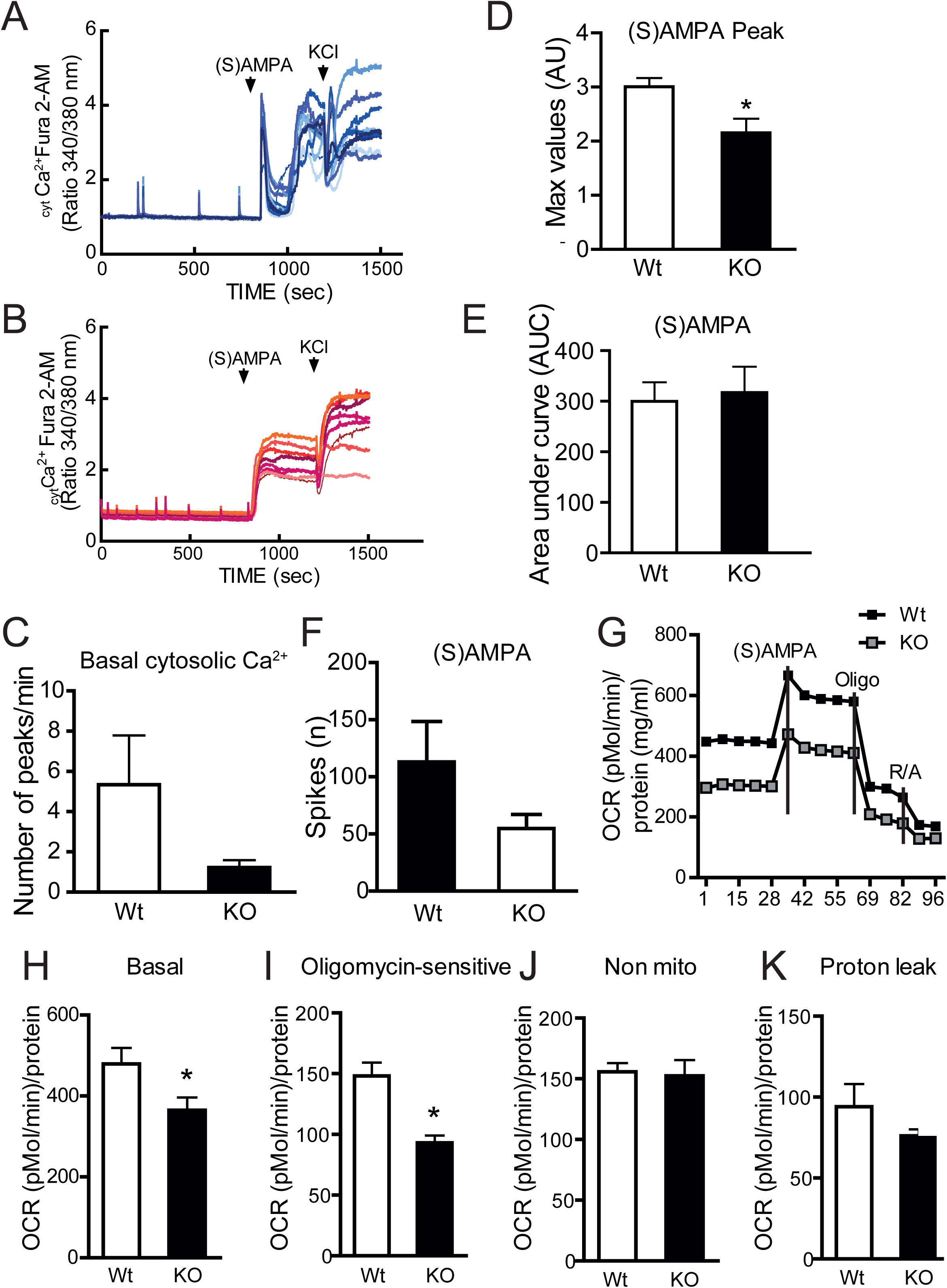
GPRIN1 physiological function in primary neurons. (A) Representative traces of Fura-2AM measurements recorded from primary cortical GPRIN1 WT and (B) GPRIN1^−/−^ neurons. Compounds application is indicated by bars above the traces. (C) The number of mini-peaks per minute was quantified during the first 5 min (basal). Max Peak (D) and Area under Curve (E) for intracellular Ca^2+^ levels after the addition of (S)AMPA (100 μM). (F) Plotted data from MEA recordings of primary neurons treated with (S)AMPA. (G) Representative oxygen consumption rate (OCR) measurements obtained with primary cultures of DIV 9 cortical neurons. At 45 min prior the experiment, culture medium was replaced with KRBH containing glucose (5 mM) followed by sequential injections of oligomycin (ATP synthase inhibitor), glutamate (150 μM) and a mix of rotenone (complex I inhibitor) and antimycin A (complex III inhibitor) (H) Basal mitochondrial OCR was measured in the first min. (I) Oligomycin sensitive OCR was derived as the difference between basal and Oligomycin inhibited OCR. (J) Non-mitochondrial OCR was calculated as the remaining OCR after Rotenone/ Antimycin A addition. (K) OCR attributed to proton leak was calculated as the difference between OCR following Oligomycin A inhibition and OCR following Rotenone/Antimycin A inhibition. The OCR data represent Mean + SD from 8 measurements from 2 independent neuronal preparations (*p<0.05).

### GPRIN1 is part of a dynamic complex of proteins affecting neuronal function

Neuronal electrical activity increases intracellular Ca^2+^ levels and activates multiple signaling molecules and gene transcription programs known to promote synapse development (Greer and Greenberg, 2008; Kelleher and Bear, 2008). In line with previous studies, we found that prolonged (S)AMPA treatment stimulated BDNF secretion (Lauterborn et al., 2003) (**Fig.S4D**), increased synaptic gene *Homer1* (**Fig. S4E**) and slightly reduced AMPA receptor subunit *Gria1* gene expression (**Fig. S4F**) in both GPRIN1^−/−^ and WT neurons. Moreover, the protein expression levels of AMPA receptor subunit (GluR1) (**Fig. S4G**) were comparable between WT and GPRIN1^−/−^neurons. These data suggest that the functional changes observed in GPRIN1 lacking neurons might be due to a signaling defect rather than an intrinsic change in receptor expression or defective regulated secretion. These results led us to conclude that GPRIN1 affects neuronal function by regulating downstream signaling pathways activated by ionotropic or GPCR receptor stimulation. Indeed, functional properties of receptor systems have been found to depend on their dynamic interactions with several intracellular signaling molecules (Ben-Shlomo et al., 2003). By combining direct immunoprecipitation (IP) of endogenous GPRIN1 in neurons followed by liquid chromatography tandem mass spectrometry (LC-MS/MS), gene ontology (GO) annotation, and pathway map enrichment analysis, we uncovered specific proteins interacting with GPRIN1. Interestingly, GPRIN1 partners were globally enriched in cytoskeleton dynamics and in synaptic vesicles trafficking and linked to major pathways associated with actin remodeling, such as Rho GTPase and G alpha signaling (**Fig. S3C-E**), clearly supporting GPRIN1’s role in downstream signaling pathways activated by neuronal receptors.

### GPRIN1^−/−^ mice showed altered learning behavior without memory impairment

Proper Ca^2+^ homeostasis and signaling are key for maintaining a functional network and highly complex pathways in the brain, such as neuronal plasticity, learning and memory. *In vivo*, the electrical activity within a network of neurons is strictly linked with fast excitatory neurotransmission that requires dynamic changes in synaptic AMPA receptor subunits distribution and expression, affecting the strength of the postsynaptic response to glutamate (Anggono and Huganir, 2012). These forms of synaptic plasticity, named long-term potentiation (LTP) and long-term depression (LTD), involve the strengthening or weakening of synaptic transmission, respectively, and are thought to be molecular-correlated with learning and memory (Kandel, 2001; Luscher and Malenka, 2012; Malenka and Bear, 2004). In order to address whether GPRIN1 might play a part in complex signaling pathways lying behind synaptic plasticity *in vivo*, neurocognitive tests were performed in adult GPRIN1^−/−^ mice. By using the Morris Water Maze (MWM) test, a classical method to assess spatial learning and memory function (Morris, 1984), we found that the GPRIN1^−/−^ mice showed learning deficits with a significant increase in escape latencies and distance compared to WT control animals (**Fig. 4AB**). However memory skills were similar during the final probe trial between the GPRIN1^−/−^ and WT mice (**Fig. 4CD**) excluding any cognitive decline as a result of GPRIN1 depletion. To spotlight emotional learning differences, the same mice were exposed to the fear conditioning (FC) test where animals learn to predict aversive events. The FC test showed minor differences in GPRIN1^−/−^ mice where the number of freezing events, expected to increase during the context test, remained unchanged compared to WT controls (**Fig. 4E**). These overall results proved that GPRIN1 might affect learning performance in adult mice, but is not by itself sufficient to give rise to cognitive deficits further confirming that GPRIN1 is more likely part of a complex with other synaptic and cytoskeleton proteins.

**Figure 4:**
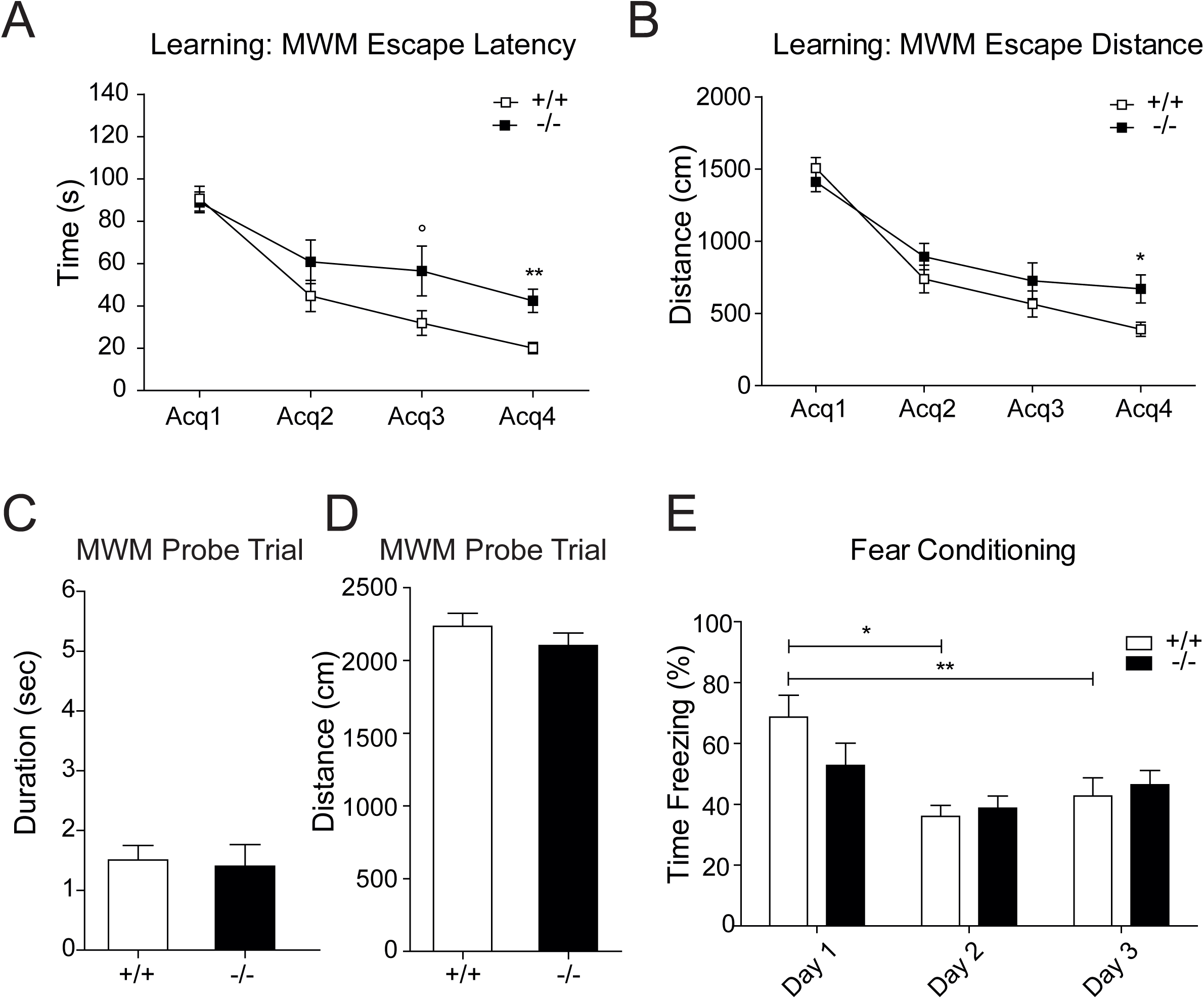
Cognitive consequences of GPRIN1 ablation in adult mice. (A) Data acquired during learning sessions of 11 GPRIN1 WT and 12 GPRIN1^−/−^ mice in MWM task showing latencies to find the escape platform and (B) radius around the escape platform and expressed as mean ± SEM. (C) Time and (D) distance traveled in the target quadrant were recorded during the final MWM probe trial. (E) Comparison and analysis of the main components of each session of the fear conditioning test (Training: day 1, Context: day 2, Tone: day 3). Percentage of time freezing for the training session (day 1), of the context session (day 2) and of the tone session (day 3) for 12 WT mice and 12 -/-mice. Data expressed as mean ± SEM.

### GPRIN1 contribution to chemical LTP (cLTP)-induced signaling mechanisms

Long-term potentiation (LTP) and long-term depression (LTD) of synaptic activity, correlating respectively with the strength or weakness of synaptic transmission, are widely used models to assess synaptic plasticity that occurs during learning and memory (Kandel, 2001; Luscher and Malenka, 2012; Malenka and Bear, 2004). Moreover cognitive performance associate with synaptic modulations (Bliss and Collingridge, 1993) (Bi and Poo, 1998) and result in a coordinated increase in synaptic signaling between neurons. This effect can be recapitulated *in vitro* via high-frequency electrical stimulation or by chemical stimulation (cLTP) and lasts from several hours up to many days (Bolshakov et al., 1997). Driven by our behavioral observations *in vivo* we opted to investigate the contribution of GPRIN1 to LTP. Potentiation of neurons without direct synaptic electrical stimulation was recorded by MEA by using a well-known pharmacological paradigm (Barad et al., 1998) (Otmakhov et al., 2004) (Oh et al., 2006). We induced cLTP by using a cocktail of forskolin (FSK), potent activator of adenylyl cyclase, and IBMX, a phosphodiesterase inhibitor to boost cyclic AMP (cAMP) production, which in turn potentiates a large fraction of synapses in the network (Otmakhov et al., 2004) and generates neuronal spikes (Niedringhaus et al., 2013). Accordingly cLTP stimulation of WT neurons increased network-wide firing rates after 3 and 24 hours. However the number of spikes recorded in GPRIN1^−/−^ neurons was dramatically reduced confirming a possible functional involvement of GPRIN1 (**Fig. 5A**). To verify changes in GPRIN1 expression and localization during high neuronal activity we performed live imaging in neurons infected with GFP-tagged GPRIN1 and challenged with cLTP. Our results showed that GPRIN1 loses its membrane-bound localization and becomes over time more prominent in the cytoplasm and along the neurites (**Fig. 5B**), suggesting a dynamic localization of GPRIN1 after receptor activation. Moreover, it has been shown that cAMP-dependent protein kinase A (PKA) phosphorylates the AMPA receptor subunit GLUR1, essential for LTP in the CA1 region of the adult hippocampus (Zamanillo et al., 1999). This specific phosphorylation event at Serine 845 (Mammen et al., 1997; Roche et al., 1996) potentiates AMPA receptor function and probably its trafficking (Lee et al., 2000). In primary neurons FSK strongly induced phosphorylation of GLUR1 independently of GPRIN1 expression (**Fig. 5C**). However, exposure to either FSK or a cAMP analog (cpt) strongly increased phosphorylated endogenous PKA targets in WT neurons but to a lesser extent in GPRIN1^−/−^ neurons (**Fig. 5D-S3B**) suggesting that during neuronal activity GPRIN1 might indirectly affect downstream effectors contributing to synaptic signal propagation.

**Figure 5:**
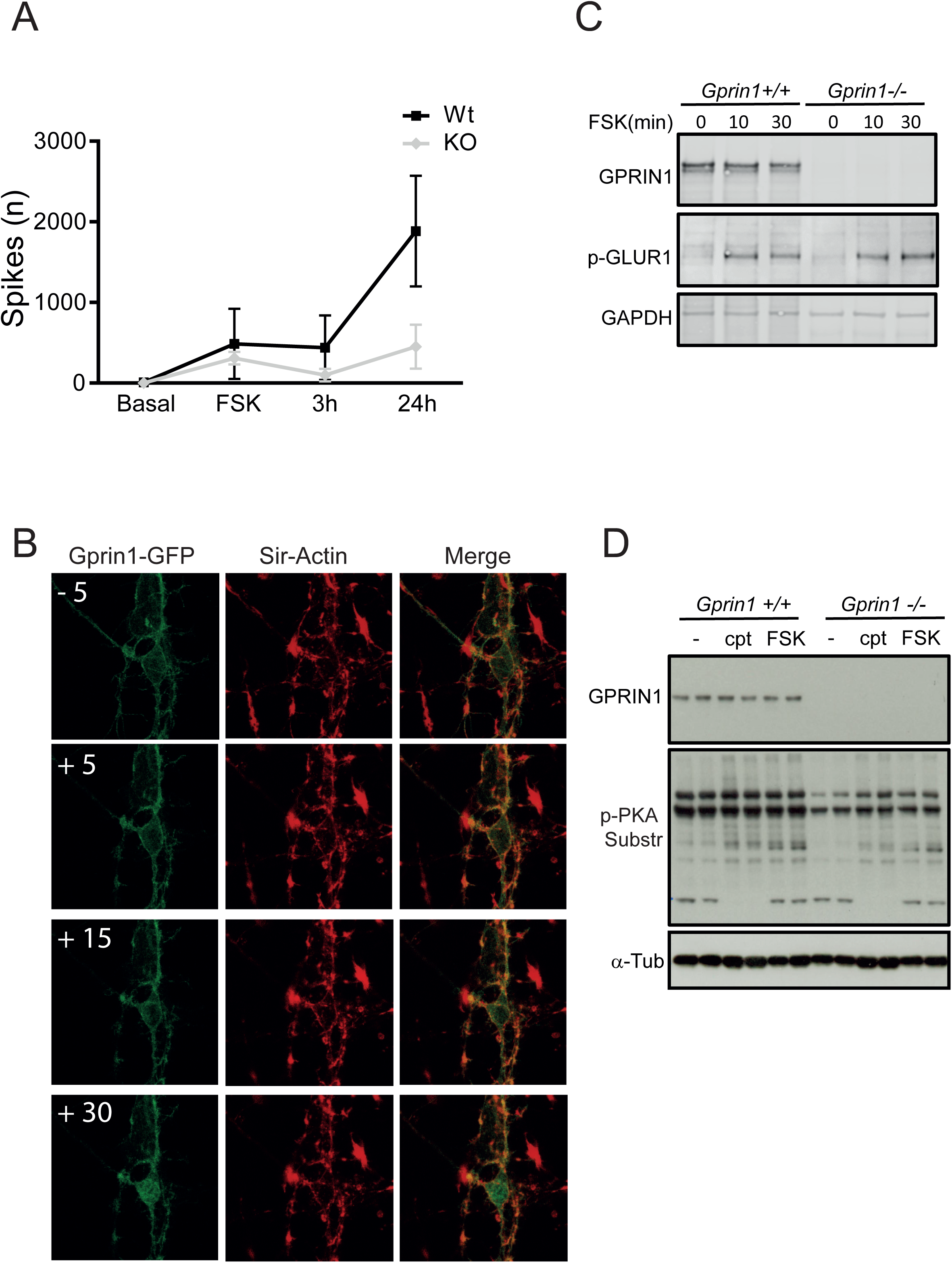
GPRIN1 role in signal transduction cascade. (A) Number of spikes recorded by MEA during a time course treatment with FSK and IBMX in GPRIN1 WT and GPRIN1^−/−^ neurons. (B) GPRIN1-GFP localization in infected GPRIN1^−/−^ neurons before and after stimulation with FSK and IBMX; Sir-Actin was used as a counterstaining to visualize the neuronal network and exclude off-target changes during the experiment. (C) Time course of FSK+IBMX stimulation showing phosphorylation of AMPA receptor subunit (GLUR1) in GPRIN1 WT and GPRIN1^−/−^ primary neurons (D) Whole lysates from GPRIN1 WT and GPRIN1^−/−^ primary neurons after 30 min of treatment with FSK (10 μM) and IBMX (100 nM) or cAMP analog cpt (10 μM) were separated by SDS PAGE and immunoblotted with the indicated antibodies.

### Lack of GPRIN1 is associated with vulnerability to neuronal stress

Among the several mechanisms involved in synaptic plasticity NMDA-type glutamate receptors (NMDARs) ensure neuronal excitability, Ca^2+^ influx and memory formation. However it is widely believed that overstimulation of NMDARs leads to excitotoxicity, characterized by disrupted Ca^2+^ homeostasis, oxidative/nitrosative stress, mitochondrial dysfunction and ultimately leading to cell death and apoptosis (Avery, 2011). The subsequent synaptic loss and dysfunction contribute to cognitive deficits in most neurodegenerative diseases, such as Alzheimer’s disease (AD) (Cornelius et al., 2013; Mattson, 2008). Given our proposed role for GPRIN1 in signal transduction downstream of receptor activation, we explored its contribution to stress response in neurons. Interestingly, long exposure of WT primary neurons to saturating concentrations of NMDA (200 μM) increased gene expression of GPRIN1 as well as of the ER stress markers Grp78/Bip (**Fig. 6AB**) however Grp78/Bip expression was significantly higher in the GPRIN1 KO neurons. Interestingly our interactome analysis showed that endogenous GPRIN1 specifically interacts with caspase-3 in primary neurons (**Fig. S3G**) as previously reported at synaptic level (Victor et al., 2018). In line with these findings we showed that GPRIN1 interacted with caspase3 in SHSY-5S neuroblastoma cell lines and AMPA stimulation likely increased this interaction (**Fig. 6C**). In order to dig deeper in the role played by GPRIN1 during ER stress and caspase 3 activation, we knocked-down endogenous GPRIN1 in SHSY-5S cells by shRNA. Our results showed that loss of GPRIN1 led to increased basal levels of caspase 3 and neuronal cells challenged with thapsigargin (TG) showed higher levels of cleaved caspase 3 (**Fig. 6D**). In addition 24 hours treatment with TG led to dramatic reduction of GPRIN1 protein levels (**Fig. 6D, Fig. S3H**). Finally, lack of GPRIN1 made neurons more sensitive to NMDA-induced stress, as demonstrated by an increase in cleaved caspase-3 (**Fig.6E**). Collectively these results strongly suggest that GPRIN1 might play a critical role in neuronal stress and apoptosis by interacting with and probably inhibiting caspase 3 activity in neurons. In conclusion, loss of GPRIN1 correlates with increased vulnerability to neuronal stress, probably linked with functional deficit during development impacting on learning behavior in adult mice.

**Figure 6:**
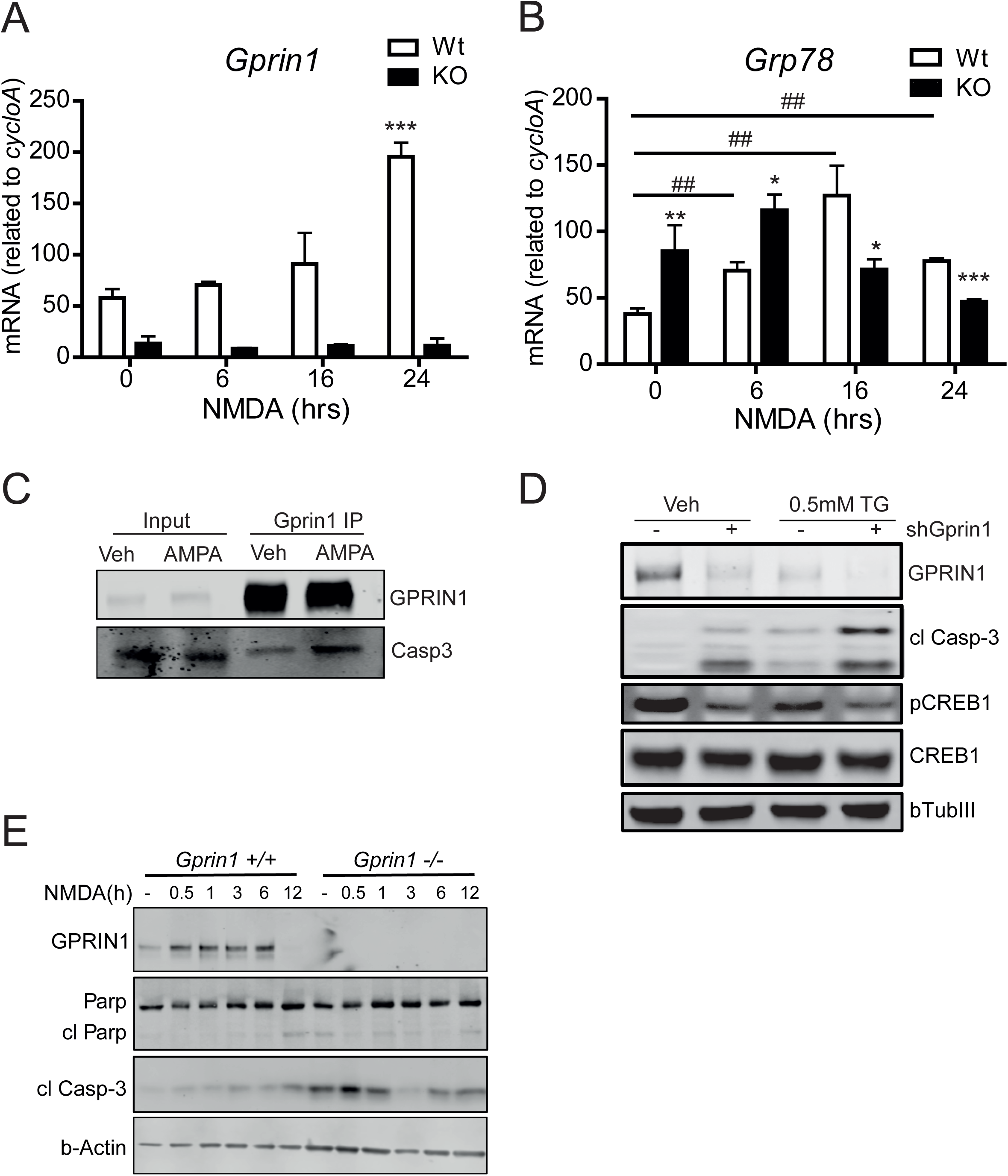
GPRIN1 in stress response and cell death. (A) qPCR analysis during time course treatment with NMDA (200 μM) showing GPRIN1 and (B) Grp78 gene expression (C) GPRIN1 was immunoprecipitated with sheep anti-GPRIN1 antibody from SHSY-5Y cells treated with vehicle (lanes 3) or (S)AMPA (lanes 4). Lanes 1-2: immunoblot analysis of cell extracts before immunoprecipitation (Input). Immunoprecipitates (IP) were analyzed by SDS-PAGE and immunoblotting (IB) using anti-caspase 3 antibody (D) WB analysis of SHSY-5Y cells treated with Thapsigargin after knock-down of endogenous GPRIN1 by shRNA (E) Whole lysates from GPRIN1 WT and GPRIN1^−/−^ primary neurons exposed to NMDA time course treatment were separated by SDS-PAGE and immunoblotted with the indicated antibodies to assess apoptotic response.

## DISCUSSION

In this study, we describe for the first time the neurological phenotypes of the GPRIN1^−/−^ mouse focusing on neuronal function and learning behavior. Despite recent advances in understanding the signaling pathway of the proteins involved in neurite outgrowth and synaptic plasticity linked to neurodevelopmental disorders, the onset of the underlying neurobiological substrates of the pathology remain largely unknown. Here we show that GPRIN1 is highly expressed in adult mouse brain, mainly enriched in the hippocampus, cortex and habenula, brain areas primarily involved in the processing of learning, memory and emotional reactions. Our results show that GPRIN1 is not essential in overall brain development but it plays a functional role both in the early stages of neuronal network staging and in fully mature neurons. GPRIN1 affects calcium signaling and receptor agonist-induced physiological neuronal activity. Our *in vivo* findings showing that GPRIN1^−/−^ mice display deficits in learning behavior without impairing memory performance further support the ex *vivo* observations. Although specific mutations of GPRIN1 have not yet been reported in correlation with human neurological or degenerative diseases, recent studies proposed GPRIN1 as a candidate target gene of Methyl-CpG binding protein 2 (MeCP2), a transcription factor encoded by an X-linked gene and mutated in a progressive neurodevelopmental disorder, called Rett syndrome (Chahrour et al., 2008). MeCP2 mutations in mice result in a neurological phenotype similar to that of human MeCP2 duplication syndrome with neurobehavioral abnormalities, ranging from mild learning disabilities to autism, X-linked mental retardation and infantile encephalopathy. Interestingly, mild and non-syndromic forms of mental retardation pathologies show little, if any, change in brain macro-anatomy, including relevant areas like the cerebral cortex and hippocampus. These findings indicate that a defective neuronal network formation and/or plasticity, rather than structural problems, might be the underlying mechanisms regulating these pathological condition diseases (Ramakers, 2002). All together, these observations support our hypothesis where GPRIN1 might affect learning performance in adult mice, but is not by itself sufficient to give rise to brain defects and cognitive alterations. Our study foresees GPRIN1 as a key player in a dynamic protein complex team involved in signal transduction pathways responsible for learning adaptation, mainly vesicle trafficking and cytoskeleton dynamics. In particular rapid remodeling of the cytoskeleton through actin-binding proteins in the postsynaptic compartment is thought to have an important function for synaptic plasticity (Rust et al., 2010) and aberrant Rho signaling seems to be key in the etiology of mental retardation (Ramakers, 2002). Notably our proteomic analysis showed novel interacting GPRIN1 partners modulating functions related to the regulation of actin organization, such as guanine nucleotide dissociation inhibitor (RhoGDI), a protein directly and indirectly associated with neurite outgrowth, cytoskeleton remodeling and spine formation (Kim et al., 2015). Furthermore these observations fit with recent unbiased proteomic screening for cannabinoid type 1 (CB1) receptor-interacting proteins in the mouse brain, directly linking GPRIN1 with the WAVE1 complex and Rac1 (Njoo et al., 2015). This dynamic protein complex is involved in the cannabinoidergic modulation of actin remodeling occurring in developing as well as mature neurons. With actin being the most abundant cytoskeletal protein found at synapses, targeting actin-binding proteins such as GPRIN1 might be of high interest for a more specific and efficient therapeutic strategy. Besides, actin remodeling had been implicated in AMPAR- and NMDAR-dependent synaptic depression (Wang et al., 2007), where availability of glutamate receptors, mainly AMPAR, at the postsynaptic membrane is strictly linked to synaptic plasticity (Sheng and Kim, 2002). In this context, we showed that lack of GPRIN1 enhanced vulnerability to NMDA-induced stress, impaired spontaneous neuronal activity and correlated with poor learning performance. It has been shown that high-frequency network burst firing or pharmacological agents that alter spontaneous network activity can affect cortical neurons against trophic deprivation– induced apoptosis. This effect is likely due to an influx of calcium through L-type voltage-dependent calcium channels activated by neuronal burst activity and depolarization directly activating intracellular pro-survival pathways (Golbs et al., 2011) (Harris et al., 2002) and inducing the expression and release of neurotrophic factors (Brigadski et al., 2005) (Kolarow et al., 2007). Therefore, GPRIN1 might play a key role in the control of mature neurons survival by ensuring spontaneous network activity patterns in developing neuronal networks. Another key event affected by GPRIN1 expression was dysregulation of Ca^2+^ dynamics and neuronal activity-dependent signaling networks known to trigger distinct gene expression responses (Bading et al., 1993; Dolmetsch et al., 2001). Among the multiple signaling molecules that modulate specific regulator factors and activate different gene transcription programs related to synapse development (Greer and Greenberg, 2008; Kelleher and Bear, 2008), we showed that GPRIN1 might directly act on downstream effectors such as PKA and CREB. Notably, most of the genes reported to be involved in neuronal activity-dependent gene transcription are strongly associated with human disorders of cognitive function (Ebert and Greenberg, 2013), further reinforcing the need to shed light on GPRIN1’s unexplored role in brain health and disease. It is worth noting that a research study related the presence of a phosphorylated form of GPRIN1 to the early stage of Alzheimer’s Disease (AD) across different rodent and human AD models (Tagawa et al., 2015). Along with this study, and given the tight interplay between immune system and inflammation in the development of AD, it is worth mentioning that GPRIN1 is also expressed in various types of immune cells. In particular, a specific function has been proposed for GPRIN1 as a scaffold protein capable of driving the interaction between α4 nAChRs and Gαi in T cells (Nordman et al., 2014) and, recently, of mediating the anti-inflammatory properties of α7 nAChRs/Gαq signaling in microglia cells (King et al., 2017). Finally a recent proteomic study identified GPRIN1 as a novel synaptic substrate for caspase 3-dependent proteolytic cleavage, the major early event in synapse loss and neurodegeneration (Victor et al., 2018). Considering our findings showed increased level of activated caspase 3 in neurons lacking GPRIN1, further investigation needs to be done to determine whether in healthy neurons GPRIN1 inhibit caspase 3 activity in neurons by sequestering it and whether GPRIN1 might be itself a proteolytic substrate of caspase 3. Currently, we report evidence linking GPRIN1 with neuronal activity-dependent signaling pathways and we suggest that GPRIN1 might be actively involved in several neuronal disorders such as autistic-like syndrome, neuroinflammation, modulation of pain and neurodegeneration.

## AUTHOR CONTRIBUTIONS

EMD and CS conceptualized the project. CS, JP, AV, CR and EH performed molecular biology and *ex vivo* studies that formed the rationale for the included studies. UDM designed the calcium experiments. EMD performed experiments in neuronal cell lines. AND and LD performed and analyzed the LC/MS data. The primary analysis of the data and the manuscript preparation was done by EMD and CS. All authors edited the manuscript and read it before submission.

## ACKNOWLEDGEMENTS

We thank the Center of PhenoGenomics (CPG) at UDP/SV EPFL for performing the behavioral studies and analysis, the Histology Core Facility and José-Luis Sanchez Garcia (NIHS) for technical support. We acknowledge Dr Andreas Wiederkehr for kindly providing helpful suggestions on the manuscript.

## SUPPLEMENTARY MATERIAL

### Interactome analysis

Immunoprecipitation (IP) was performed using specific sheep antibody against GPRIN1 bound to protein beads to specifically isolate GPRIN1 protein complex. The interacting proteins were further purified with SDS-PAGE (*i.e.*, stacking gel-like approach without performing separation) followed by in-gel tryptic digestion and liquid chromatography tandem mass spectrometry (LC-MS/MS) for protein identification. Gene annotation and pathway enrichment analysis was performed using the MetaCore software (GeneGo, Thomson Reuters). Carbachol was used at final concentration of 100 μM.

### Human iPSC-derived (iCell) neurons

Neurons derived from human-induced pluripotent stem cells (iCell Neurons) were purchased from Cellular Dynamics International (Madison, WI, USA). Neurons were maintained according to the manufacturer’s protocol. Depending on the experiments, 140,000 cells/well were seeded on 6well plates or MatTek coverslips double-coated with 0.01% Poly-L-Ornithine and 3.3 μg/ml laminin (Sigma-Aldrich; St. Louis, MO, USA).

## SUPPLEMENTARY FIGURE LEGENDS

### Supplementary Figure 1

(A) IF Hippocampus WT/KO: GPRIN1 and Dapi

(B) HE staining (WT/KO) hippocampus

(C) IF Habenula WT: GPRIN1 and Dapi

(D) IF Hypothalamus WT: GPRIN1 and Dapi

(E) IHC Hippocampus WT: GPRIN1, Gap43 and Dapi

(F) IHC Hippocampus WT: GPRIN1, S100 and Dapi

(G) Synaptic marker Snap25 mRNA expression (qPCR)

(H) Gene expression of several neuronal markers in hippocampal neurons (WT/KO)

### Supplementary Figure 2

(A) iCell neurons were fixed and stained with Dapi (Invitrogen), GPRIN1 and specific neuronal marker (bTubIII)

(B) Higher-magnification image depicting the punctate GPRIN1 distribution along a single neurite

(C) Human iPSC-derived neurons (IF) GPRIN1, GAP43 and Dapi

(D) Human iPSC-derived neurons expression markers (qPCR)

### Supplementary Figure 3

(A) Area under curve (AUC) of intracellular calcium levels measured with Fura-2AM after the addition of Carbachol (100 μM)

(B) Phosphorylation of downstream PKA-substrates in neurons after 30 min of treatment with FSK+IBMX or cpt by immunoblotting with indicated antibodies.

(C) IP of GPRIN1 on primary neurons WT/KO (enrichment control with SDS-PAGE before LC-MS/MS analysis)

(D) Canonical pathway map enrichment analysis on GPRIN1 interactors

(E) GO process networks enrichment analysis on GPRIN1 interactors

(F) GO localization enrichment analysis on GPRIN1 interactors

(G) List of proteins (identified with LC-MS/MS after IP of GRIN1) potentially interacting with GPRIN1

(H) Whole lysates from SHSY-5Y cells treated with a dose response of Thapsigargin were separated with SDS-PAGE and immunoblotted with the indicated antibodies

### Supplementary Figure 4

(A) Schematic explaining the Seahorse XF Extracellular Flux 96 analyzer. The standardized and sequential addition of respiratory chain inhibitors allows investigating several domains of mitochondrial respiration.

(B) AMPA representative

(C) Quantification of oxygen consumption rate (OCR) after (S)AMPA, Glutamate and KCl stimulation

(D) Secretion of BDNF detected by ELISA in culture supernatants from neurons treated with (S)AMPA (100 μM) for 5 and 10 min.

(E) qPCR analysis of AMPA receptor subunit Gria1 in GPRIN1 WT and GPRIN1^−/−^ primary cortical neurons treated ON (16 hours) with (S)AMPA 100 μM

(F) qPCR analysis of synaptic marker Homer1 mRNA expression in GPRIN1 WT and GPRIN1^−/−^ primary cortical neurons treated ON (16 hours) with (S)AMPA 100 μM.

(G) Whole lysates from GPRIN1 WT and GPRIN1^−/−^ primary neurons were separated by SDS PAGE and immunoblotted with the indicated antibodies

---

**Supplementary Table 1:**
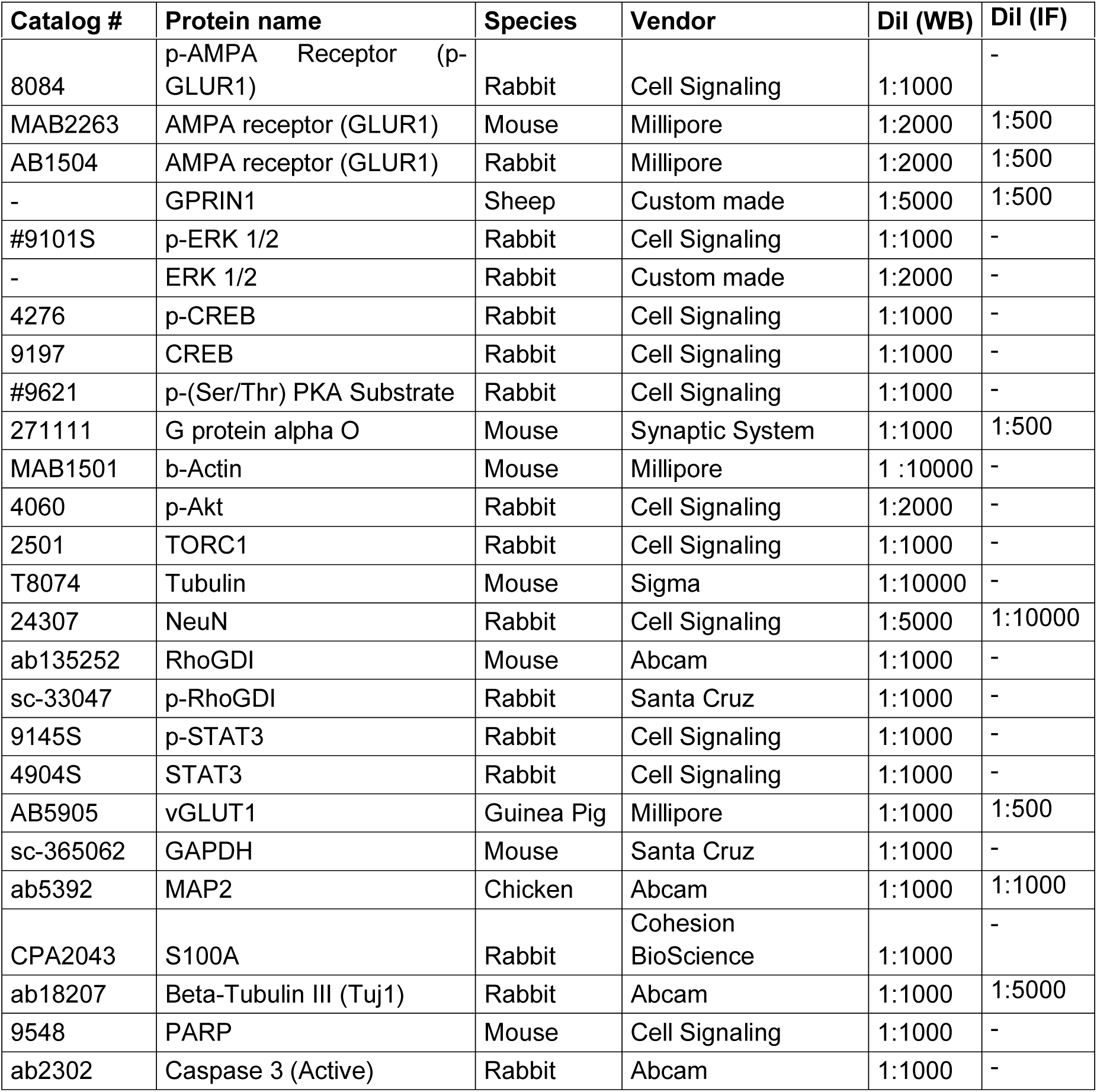

**Supplementary Table 2:**
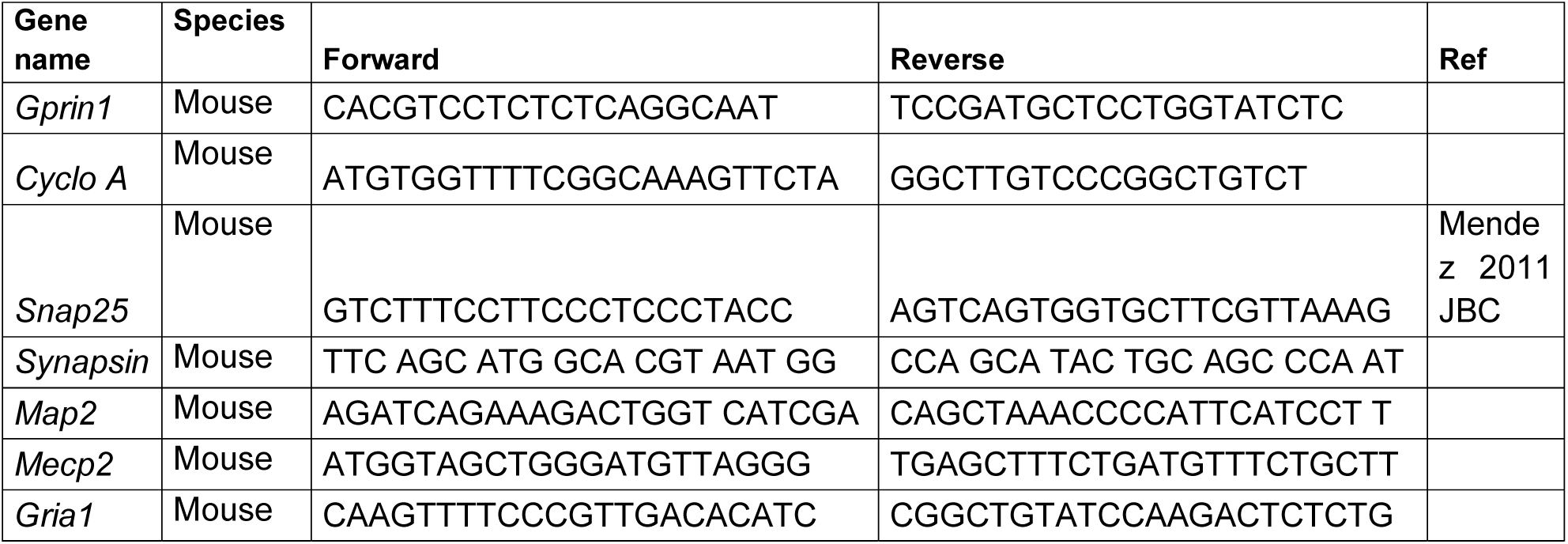
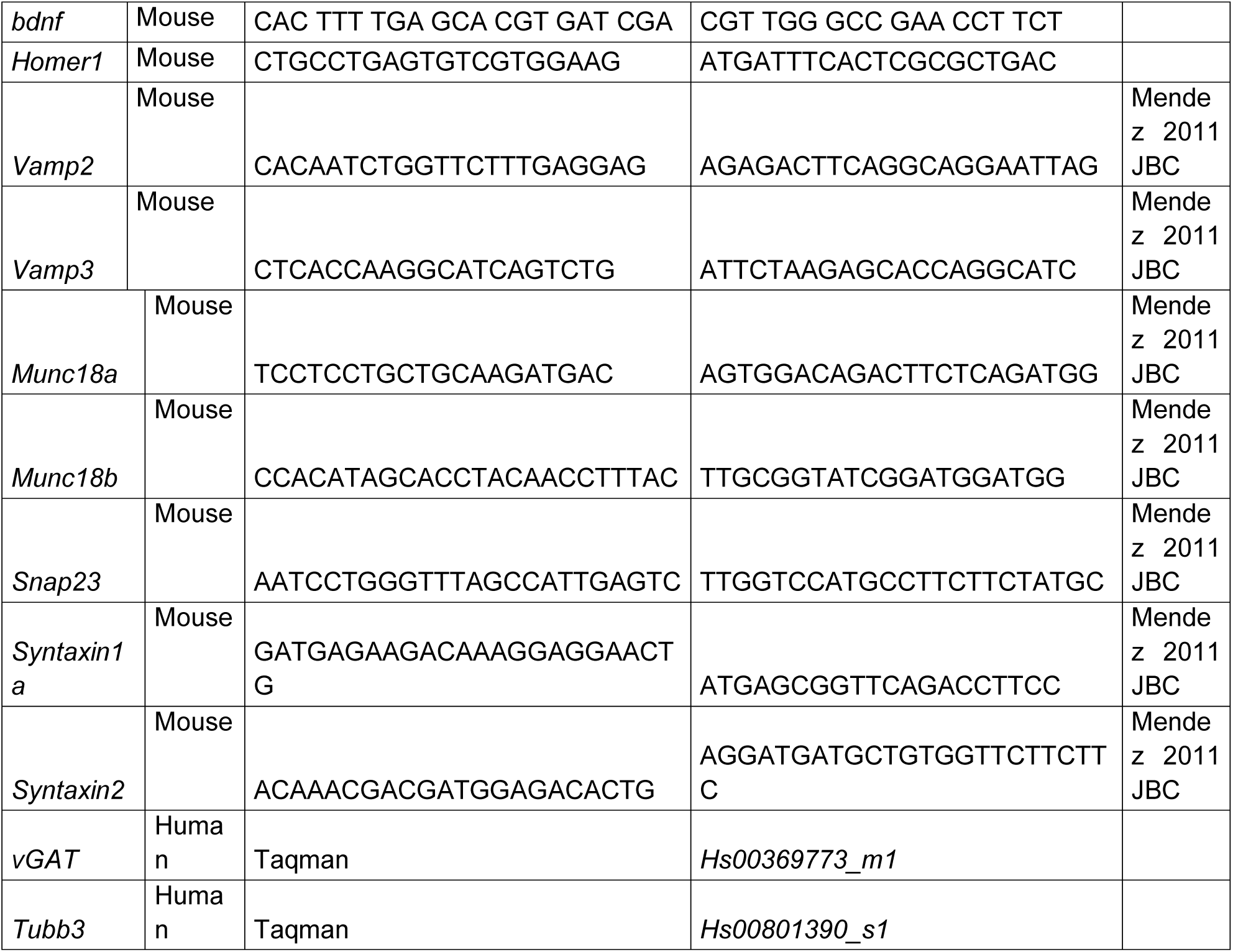

